# The subunit 3 of the SUPERKILLER (SKI) complex mediates miR172-directed cleavage of *Nodule Number Control 1* (*NNC1*) to modulate nodulation in *Medicago* truncatula

**DOI:** 10.1101/2025.01.06.631520

**Authors:** Soledad Traubenik, Mauricio Alberto Reynoso, Francisco Sánchez-Rodríguez, Milagros Yacullo, Aurelie Christ, Maureen Hummel, Thomas Blein, Martín Crespi, Julia Bailey-Serres, Flavio Antonio Blanco, María Eugenia Zanetti

## Abstract

Legumes and rhizobia establish a nitrogen-fixing symbiosis that involves the formation of a lateral root organ, the nodule, and the infection process that allows intracellular accommodation of rhizobia within nodule cells. This process involves significant gene expression changes regulated at the transcriptional and post-transcriptional levels. We have previously shown that a transcript encoding the subunit 3 of the Superkiller Complex (SKI), which guides mRNAs to the exosome for 3′-to-5′ degradation, is required for nodule formation and bacterial persistence within the nodule, as well as the induction of early nodulation genes (e.g., *MtENOD40*) during the *Medicago truncatula*-*Sinorhizobium meliloti* symbiosis. Here, we reveal through transcript degradome and small RNA sequencing analysis that knockdown of *MtSKI3* impairs the miR172-directed endonucleolytic cleavage of the mRNA encoding Nodule Number Control 1 (MtNNC1), an APETALA2 transcription factor that negatively modulates nodulation. Knockdown of *MtNNC1* enhances nodule number, bacterial infection, and the induction of *MtENOD40* upon inoculation with *S. meliloti* whereas overexpression of a miR172-resistant form of *MtNNC1* significantly reduces nodule formation. This work identifies miR172 cleavage of *MtNNC1* and its control by MtSKI3, a component of the 3′-to-5′mRNA degradation pathway, as a new regulatory hub controlling indeterminate nodulation.

## Introduction

mRNA decay is a key regulatory process that modulates gene expression by affecting the stability and, thus, the steady-state levels of cellular mRNAs for protein synthesis. Decay of mRNAs occurs either through the action of exoribonucleases or the endonucleolytic cleavage guided by microRNAs (miRNAs) and small interference RNAs (siRNAs). Bulk degradation of cytoplasmic mRNAs by exonucleases is accomplished by distinct deadenylation-dependent and deadenylation-independent pathways (Chantarachot and Bailey-Serres, 2018). In the deadenylation-dependent pathways, the protecting polyA tail at the 3′ end of translationally repressed mRNAs is removed by one of the different types of deadenylases. Subsequently, the deadenylated mRNAs can undergo decapping and 5′-to-3′ decay, which in plants is catalyzed mainly by the 5′-to-3′exoribonuclease XRN4 (Potuschak et al., 2006; Rymarquis et al., 2011) or be guided by the SUPERKILLER (SKI) complex to the RNA exosome, where degradation is in the 3′-to-5′ direction (Liu and Chen, 2016; Lange and Gagliardi, 2021). The deadenylation-independent pathway occurs co-translationally in the 5′-to-3′ direction on ribosome-associated mRNAs, whose translation has been paused or stacked at upstream open reading frames (uORFs), the stop codon of main open reading frames (mORFs). or in the proximity of non-cleavable miRNA target sites (Yu et al., 2015; Hou et al., 2016; Yu et al., 2016). This pathway, which requires the removal of the 5′cap and the function of XRN4, exhibits a decay with a three-nucleotide periodicity that reflects the codon-by-codon translocation of elongating ribosomes along the mRNA (Yu et al., 2015; Yu et al., 2016; Pelechano et al., 2015; Carpentier et al., 2020). Degradation of mRNAs by endonucleolytic cleavage is guided by miRNAs or siRNAs bound to a member of the ARGONAUTE (AGO) protein family in a process known as post-transcriptional gene silencing (PTGS) (Bartel, 2004; Vaucheret, 2006). mRNA endonucleolytic cleavage results in the production of a deadenylated 5′cleavage fragment and a decapped 3′ cleavage fragment, which can subsequently be degraded by the SKI/RNA exosome complex and XRN4, respectively. Also, both 5′ and 3′ fragments can be substrates of RNA-dependent RNA polymerase 6 (RDR6), triggering the production of secondary siRNAs (Allen et al., 2005; Yoshikawa et al., 2005; Manavella et al., 2012).

SKI is an exosome-associated complex evolutionary conserved in eukaryotes with key functions in the deadenylation-dependent 3′-to-5′ mRNA degradation, the surveillance mechanisms of nonsense-mediated decay (NMD) and non-stop decay (NSD) (Arribere and Fire, 2018; Szádeczky-Kardoss et al., 2018), as well as in small RNA-mediated endonucleolytic cleavage of mRNAs in metazoa and plants (Orban and Izaurralde, 2005; Zhang et al., 2015). The SKI complex is a tetramer composed of SKI2, SKI3, and two SKI8 subunits. The SKI2 subunit functions as a helicase that unwinds the RNA, the SKI3 subunit is a scaffold protein with tetratricopeptide repeats (TPR), whereas SKI8 is a WD40 repeat-containing protein. Both TPR and WD40 repeats have been involved in protein-protein interactions (Halbach et al., 2013). In animals, different lines of evidence suggest that the SKI complex can act co-translationally to channel translationally stalled mRNAs to the RNA exosome for degradation through its association with 80S ribosome-bound mRNAs (Schmidt et al., 2016; Zinoviev et al., 2020; Kögel et al., 2022). In yeast, biochemical and cryo-electron microscopy (EM) structural analyses have shown that the SKI complex binds the 80S ribosome via SKI2, the N-terminal of SKI3, and one of the SKI8 subunit (Schmidt et al., 2016), whereas in mammals the interaction of the SKI complex with the 40S ribosomal subunit is exclusively mediated by SKI2 encompassing a gatekeeping mechanism that switches from a closed conformational state to an ATP induced open conformational state that facilitates the extraction of 80S ribosome-bound mRNAs and their delivery to the RNA exosome for degradation (Kögel et al., 2022). In plants, the interaction of the SKI complex with 80S ribosomes and any role in the co-translational RNA surveillance pathway remains unexplored. Instead, genetic and molecular evidence points to the role of the SKI complex in preventing the ectopic production of secondary siRNA from endogenous miRNA targets and transgenes in Arabidopsis (Branscheid et al., 2015; Yu et al., 2015; Zhang et al., 2015; Vigh et al., 2022). Simultaneous disruption of bidirectional RNA decay (i.e., 5′-to-3′and 3′-to-5′) by mutations in genes encoding subunits of the SKI complex and the cytoplasmic 5′-to-3′ exoribonuclease EIN5, enhanced the production of 21 and 22 siRNAs in Arabidopsis, indicating that 3′-5′ and 5′-3′ RNA decay pathways mediated by the SKI complex and EIN5, respectively, suppress post-transcriptional gene silencing (PTGS) (Zhang et al., 2015). Concomitantly, Yu et al (2015) reported that mutation of *SKI3* restores sense PTGS in a hypermorphic *ago-1* mutant background, whereas Branscheid et al. (2015) reported that mutation of Arabidopsis *SKI2* enhanced the production of secondary siRNAs generated from either the 5′ or the 3′ cleavage fragments of miRNA targets and these siRNAs mapped close the miRNA cleavage site. Also, the authors found that mutation in any of the *SKI* genes, *SKI2*, *SKI3,* or *SKI8*, compromised degradation of the 5′-cleavage fragments, but not 3′-cleavage fragments, derived from miRNA endonucleolytic cleavage (Branscheid et al., 2015). A subsequent study revealed that in the absence of a functional SKI complex, most of the transcripts that accumulate in the cytoplasm are redirected to the 5′-to-3′ XRN4 pathway for degradation (Zhao and Kunst, 2016). More recently, Vigh et al, (2022) showed that inactivation of the exosome subunit RRP45B results in the accumulation of 5′-cleavage fragments and enhanced production of secondary siRNAs, similar to that observed in *ski2* mutants. Interestingly, mutation of the Release Factor PELOTA1, which is required for ribosomal subunit dissociation of stalled ribosome, led to the production of siRNAs from miRNA targets that overlap but are distinct from those produced in the *ski* and *rrpb45b* mutants, suggesting that PELOTA limits siRNA amplification by reducing ribosome stalling (Vigh et al., 2022).

In a previous study, we identified a transcript encoding the SKI3 subunit in the model legume *Medicago truncatula* (*MtSKI3*), which is subjected to translational regulation during the nitrogen-fixing symbiosis with *Sinorhizobium meliloti* (Traubenik et al., 2020). Knockdown of *MtSKI3* results in a reduction in the number of nitrogen-fixing specialized organs, i.e., the nodules, but also prevents the persistence of the bacteria within these nodules, compromising nodule viability and nitrogen fixation (Traubenik et al., 2020). Considering that disruption of one mRNA degradation pathway might alter the functioning of other degradation pathways, in this study we investigated whether knockdown of *MtSKI3* altered the 5′-to-3′degradation and/or miRNA guided-endonucleolytic cleavage of mRNAs in *M. truncatula* roots under symbiotic and non-symbiotic conditions by use of the genome-wide mapping of uncapped and cleaved transcripts (GMUCT) (Willmann et al., 2014). GMUCT identifies the 3’ cleavage products of miRNA or siRNA guided-endonucleolytic cleavage and other uncapped mRNAs by selecting those molecules with a free 5′ monophosphate (5′P). Hundreds of genes with free 5′P reads enriched or depleted in *MtSKI3* silenced roots were identified. One of the 3′ cleavage fragments with lower abundance in *MtSKI3* silenced roots was the product of endonucleolytic cleavage mediated by miR172 on its target mRNA, which encodes an AP2 transcription factor referred to as Nodule Number Control 1 (NNC1). In soybean (*Glycine max*), GmNNC1 acts as a negative modulator of the formation of determinate nodules and a transcriptional repressor of the *early nodulin 40* (*GmENOD40*) gene (Wang et al., 2014; Wang et al., 2019). Here, we show that whereas *MtNNC1* mRNA levels decrease upon inoculation with *S. meliloti* in control plants, they remain high in *MtSKI3* silenced roots. The silencing of *MtSKI3* does not alter the abundance of miR172, rather it leads to impaired endonucleolytic cleavage of the *MtNNC1* transcript. Knockdown of *MtNNC1* leads to more numerous nodules and infection events, as well as higher levels of *MtENOD40*, whereas overexpression of a miR172-resistant form of *MtNNC1* recapitulates the reduced nodulation phenotype observed in *MtSKI3* silenced roots (Traubenik *et al*., 2020). These results uncover a role of the SKI complex in the cleavage of *MtNNC1* mediated by miR172 during the establishment of the root nodule symbiosis.

## Results

### *SKI3*-dependent RNA fragments with a free 5′P accumulate in mock and symbiotic conditions

The underlying mechanisms by which MtSKI3 mediates nodule formation and bacterial viability are unknown. To evaluate whether RNA interference (RNAi) of *MtSKI3* in *M. truncatula* roots is altering the 5′to 3′ RNA decay and/or sRNA-guided endonucleolytic cleavage in response to rhizobia, we performed a degradome analysis by applying the GMUCT 2.0 method (Willmann et al., 2014) on *GUS* RNAi and *SKI3* RNAi roots inoculated with *S. meliloti* or with water (mock). This approach specifically identifies products of miRNA or siRNA guided-endonucleolytic cleavage and decapped mRNAs by selecting those molecules with a free 5′P (Figure 1A). Hundreds of 5′P end peaks differentially accumulate in *SKI3* RNAi roots as compared to *GUS* RNAi roots under symbiotic (94 up and 154 down) and mock conditions (118 up and 59 down) with 2<FC<0.5 and *p*-value <0.05 (Figure 1B, Supplemental Dataset 1). Under mock conditions, transcripts with increased 5′P end peaks in *SKI3* RNAi roots versus *GUS* RNAi roots are enriched in the functional categories of Perception/Signaling and Metabolism, whereas transcripts with decreased 5′P peaks are enriched in the categories of Transcriptional Regulation and Perception/Signaling (Supplemental Figure 1A, Supplemental Dataset 2). Similarly, under symbiotic conditions, the prevalent functional categories in *SKI3* RNAi roots versus GUS RNAi are Transcriptional Regulation, Metabolism, Perception/Signaling, and Chromatin remodeling (Supplemental Figure 1B). Interestingly, when comparing genes with differential 5′end peaks in *S. meliloti* inoculated versus mock roots functional categories are more diverse in *SKI3* RNAi roots than in *GUS* RNAi, including Transcriptional Regulation and Perception/Signaling as well as Cell Wall Remodeling and DNA/RNA Metabolism (Supplemental Figure 1C and 1 D, Supplemental Dataset 2). In the category of Transcriptional Regulation, transcription factors encoded by transcripts with differential 5′P peaks belong to diverse gene families, including the Auxin Response Factor (ARFs), APETALA2/Ethylene Response Factor (AP2/ERF), the basic Helip-Loop-Helix (bHLH), the basic helix loop helix (bHLH), the C2C2-Dof, the zinc finger C2H2 (Cys_2_His_2_), the Homeobox WOX, MYB and the WD40 repeats families (Supplemental Figure 2).

**Figure 1.**
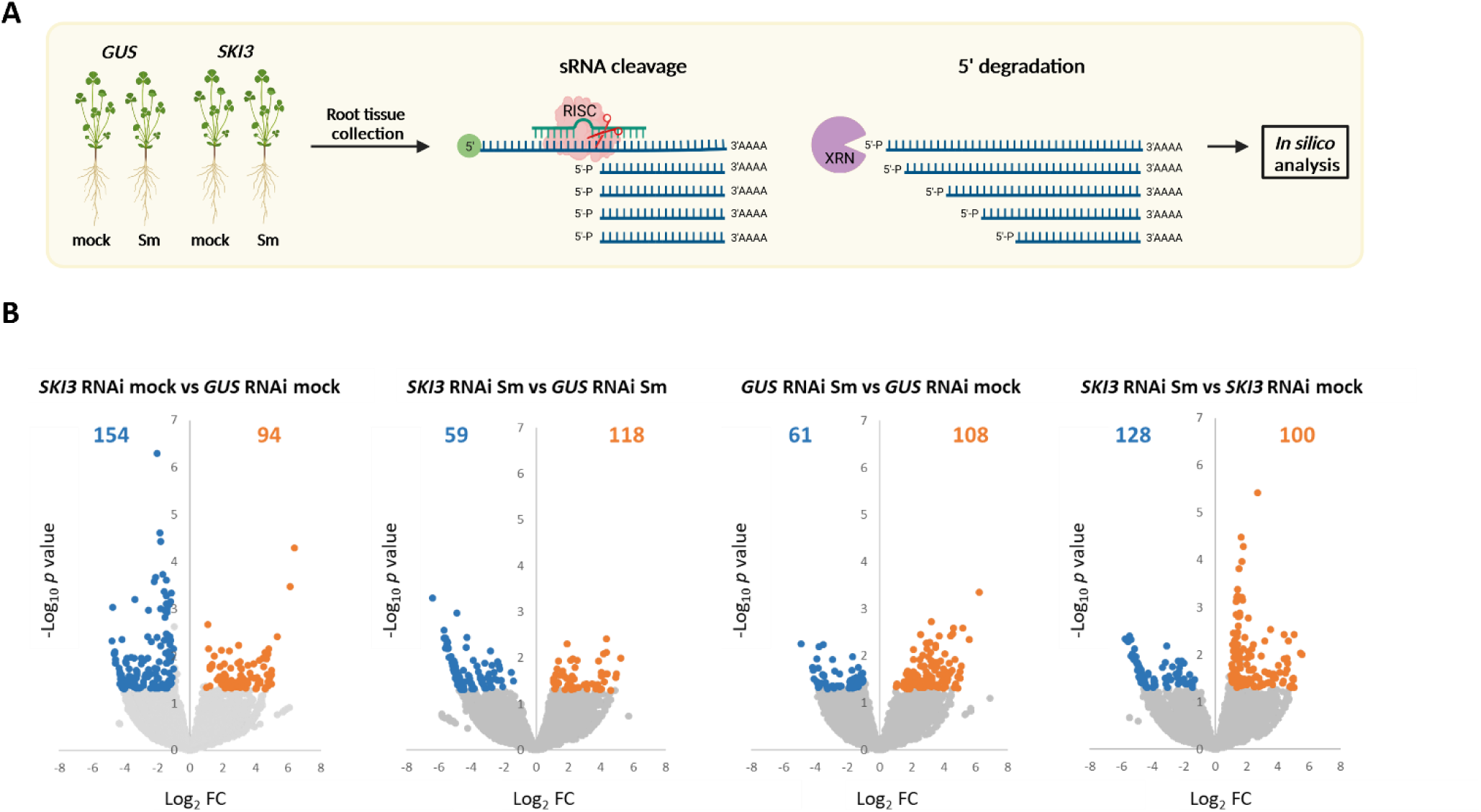
Transcripts with differential free 5′P peaks in mock and *S. melilot* inoculated *GUS* RNAi and *SKI3* RNAi roots. **(A)** Schematic overview of the experimental design using GMUCT to identify transcripts with 5′P. Roots of *M. truncatula* were inoculated with water (mock) or with *S. meliloti* (Sm). Total RNA was used to generate degradome libraries of RNA molecules with a free 5′P caused by decapping and 5′-to-3′ ′degradation by XRN4 or by sRNA-mediated endonucleolytic cleavage. Created with Biorender.com. (**B)** Volcano plots representing changes in the abundance of transcripts with a free 5′P peak. Each dot represents one transcript. Gray dots represent transcripts with no significantly different 5′P peaks in each comparison, blue dots represent transcripts with significantly downregulated 5′P peaks (FC<0.05, *p*-value <0.05) and orange dots represent transcripts with significantly upregulated 5′P peaks (FC>2, *p*-value <0.05. Numbers in each plot indicate the number of upregulated and downregulated transcripts with 5′P peaks.

### *SKI3-*dependant miRNA-directed target cleavage of specific transcripts

We then investigated whether miRNA-mediated cleavage is affected by the silencing of *MtSKI3*. We predicted the targets and their cleavage sites for the conserved canonical miRNAs of *M. truncatula* annotated in miRBase (Kozomara and Griffiths-Jones, 2014) using the miRanda algorithm (Betel et al., 2010) and then determined whether there was evidence of cleavage at these targets using degradome data allowing a window of ± 2 nucleotides from the predicted cleavage site. Our analysis confirmed 615 miRNA targets containing 5′P ends cleavage reads at the predicted miRNA cleavage site (Supplemental Dataset 3) indicating that our degradome data can robustly quantify 5′P end reads produced by miRNA-mediated cleavage. Degradome data was analyzed to quantify products of miRNA guided-endonucleolytic cleavage in both *GUS* RNAi and *SKI3* RNAi roots under mock and symbiotic conditions, and differential 5′P peaks in all pairwise comparisons were identified using DEseq2 using a *p*-value<0.05 (Love et al., 2014) (Figure 2A). Under mock conditions, we identified five transcripts with significant downregulated 5′P peaks in *SKI3* RNAi roots as compared to *GUS* RNAi encoding the Auxin Signaling F-Box 3 receptor (MtAFB3), the E3 ubiquitin-protein ligase (MtBRE1), a heterogeneous nuclear ribonucleoprotein (MtRNP), a CBL interacting protein kinases (MtCIPK6) and an ethylene-responsive transcription factor of the AP2 family, which was designated as MtNNC1 (see below). Only a single transcript showed upregulated 5′P peaks in *SKI3* RNAi roots as compared to *GUS* RNAi, encoding an E6-like protein. Under symbiotic conditions, two transcripts showed differential accumulation of 5′P peaks in *SKI3* RNAi as compared with *GUS* RNAi roots, one upregulated transcript encoding the eukaryotic translation initiation factor 5B (MteIF5B), and one downregulated transcript encoding MtAFB3. These results suggest that silencing of *MtSKI3* does not globally enhance miRNA endonucleolytic cleavage of *M. truncatula* targets, but rather affects the miRNA-mediated cleavage of specific miRNA targets, mainly reducing the amount of 3′cleavage products with a free 5′P derived from miRNA cleavage. In *GUS* RNAi roots, inoculation with *S. meliloti* diminished the degradation of only one transcript encoding a proline-rich extensin-like protein (EPR1). Conversely, in *SKI3* RNAi roots, the transcript encoding the E6-like protein exhibited 5′P fragments downregulated in response to *S. meliloti*, whereas three transcripts showed the opposite behavior. These transcripts encode an Extensin domain-containing protein (MtEXT3), the pre-mRNA-splicing factor (MtPRP38), and a hypothetical protein (MtrunA17Chr4g0.040901). Two of the transcripts with down-regulated 5′P peaks in *SKI3* RNAi versus *GUS* RNAi roots were targets of evolutionary conserved miRNAs, i.e., the transcripts encoding the AP2 transcription factor MtNNC1 and MtAFB3 are targeted by miR172 and miR393, respectively (Figure 2B and 2C, respectively). Both miRNAs and their target mRNAs have been shown to play a role during nodulation in soybean plants (Wang et al., 2014; Cai et al., 2017; Wang et al., 2019).

**Figure 2.**
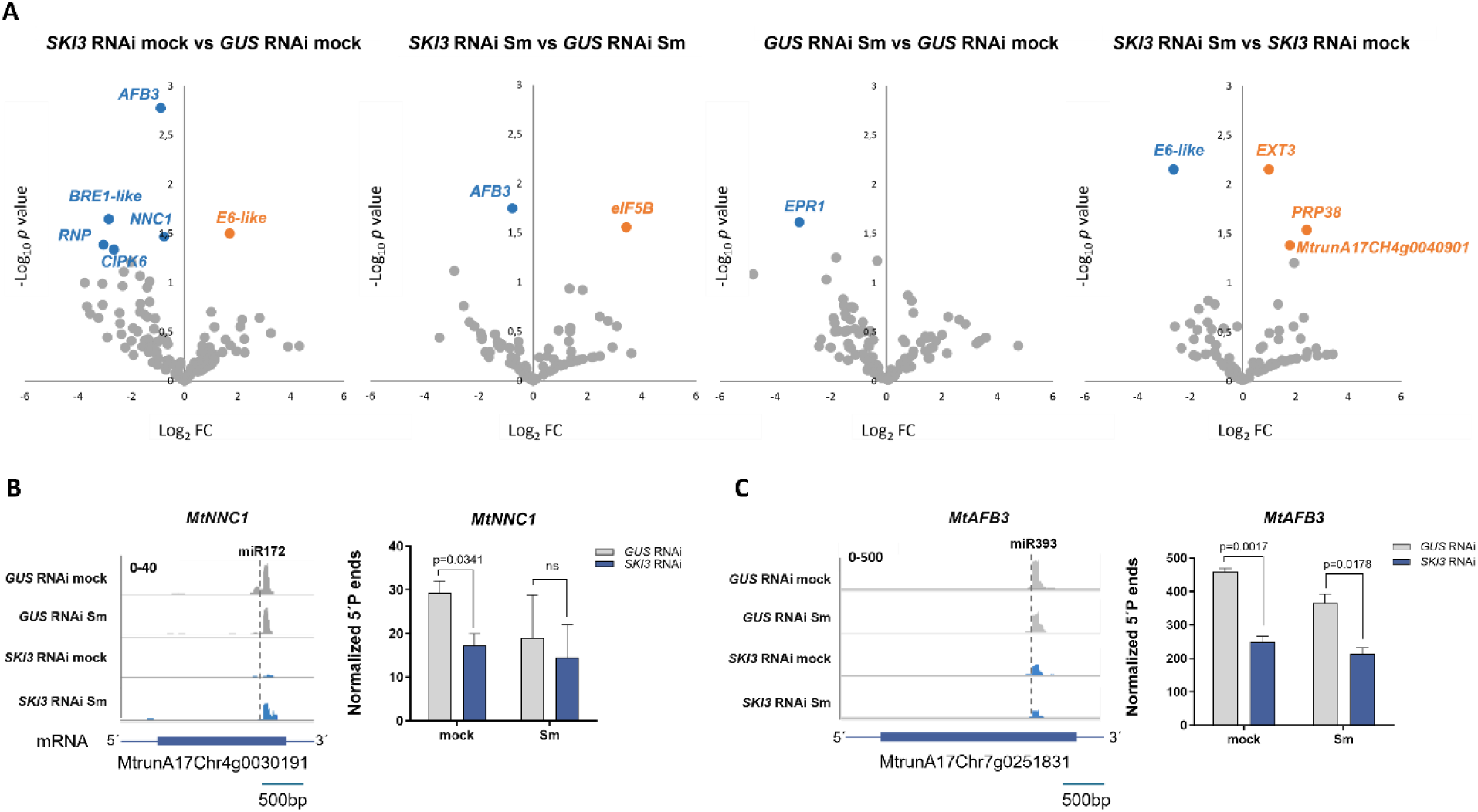
miRNA target transcripts with differential free 5′P peaks in mock and *S. meliloti* inoculated *GUS* RNAi and *SKI3* RNAi roots. (**A**) Volcano plots representing miRNA target transcripts that contain a free 5′P. Each dot represents one transcript. Gray dots represent transcripts with no significantly different 5′P peaks in each comparison, blue dots represent transcripts with significantly downregulated 5′P peaks (FC<0.58, *p*-value <0.05, and orange dots represent transcripts with significantly upregulated 5′P peaks (1.7<FC, *p*-value <0.05). (**B-C**) 5’P read coverage of *MtNNC1* (**B**) and *MtAFB3* (**C**). GUS RNAi samples are shown in gray, and *SKI3* RNAi samples in blue. Left panels: The dashed line indicates the cleavage sites of miR172 (**B**) and miR393 (**C**). Transcript models are displayed at the bottom. The numbers on the top left indicate the maximum read value of the scale, which was the same for all samples. Right panels: Normalized counts of 5′P reads ends obtained from GMUCT experiments in *SKI3* RNAi and *GUS* RNAi roots inoculated with *S. meliloti* (Sm) or water (mock). *p*-values obtained by DEseq2 for each comparison are indicated. ns: no significant difference.

### Downregulation of *MtNNC1* mRNA levels in response to rhizobia is impaired by silencing of MtSKI3

The gene encoding the AP2 transcription factor identified here (MtrunA17Chr4g0030191) is the best homolog and syntenic to *Glycine max Nodule Number Control 1* (*NNC1*) (Figure 3A), which has been described as a negative regulator of nodule initiation (Wang et al., 2014) and later, involved autoregulation of nodule number (Wang et al, 2019); thus, we named it as *MtNNC1*. To further link the action of *MtSKI3* to symbiotic nodulation, we focused our analysis on this miRNA target, since its regulation by MtSKI3 might help to explain the impaired nodulation phenotype previously observed in *SKI3* RNAi roots (Traubenik et al., 2020). The reduced number of 5′P end reads found in the *MtNNC1* transcript in *SKI3* RNAi roots as compared to *GUS* RNAi roots suggested that cleavage by miR172 might be compromised in *SKI3* silenced roots. Thus, we analyzed whether *MtNNC1* mRNA levels were modified by silencing of *MtSKI3* under mock and symbiotic conditions using RT-qPCR and primers that span at each side of the miR172a cleavage site. We found that levels of non-cleaved *MtNNC1* were significantly downregulated at 2 and 10 dpi with *S. meliloti* in *GUS* RNAi roots, in agreement with the previously reported repression in soybean roots under symbiotic conditions (Wang et al., 2014). However, this repression in response to rhizobia was abolished in *SKI3* RNAi roots at all the time points analyzed, indicating that inactivation of the SKI complex compromises rhizobia-induced miR172 mediated repression of *MtNNC1* (Figure 3B).

**Figure 3.**
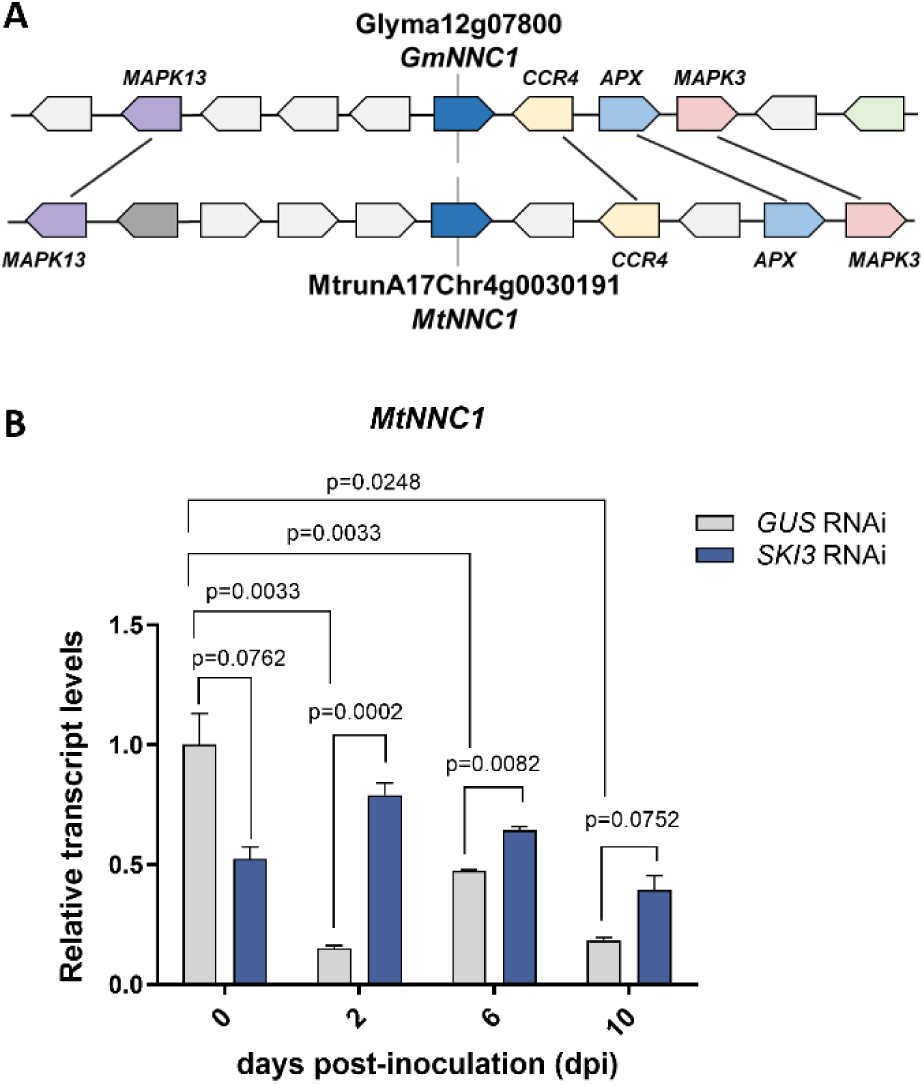
Synteny and expression analysis of *MtNNC1* in *GUS* RNAi and *SKI3* RNAi roots during symbiosis. **(A)** Syntenic regions of *M. truncatula* and *G. max NNC1*. Anchor genes flanking *MtNNC1* and *GmNNC1* are connected with colored lines. **(B)** Relative transcript levels of *MtNNC1* in *GUS* RNAi and *SKI3* RNAi roots inoculated with water (0 dpi) or with *S. meliloti* at 2, 6, and 10 dpi. Expression values were determined by RT-qPCR, normalized to *MtHIS3L*, and plotted relative to the mock sample at time 0. Each bar represents the mean ± SEM of three biological replicates. *p* values in an unpaired two-tailed Student’s *t*-test are indicated for each comparison.

### *MtNNC1* showed translational downregulation during symbiosis

*MtSKI3* was previously found to be upregulated at the translational level at early stages (2 dpi with *S. meliloti*) of the root nodule symbiosis (Traubenik et al., 2020). Since miR172 was shown to act at the level of translational repression in Arabidopsis plants (Aukerman and Sakai, 2003; Chen, 2004) and we previously found that miR172 was bound to translating ribosomes in *M. truncatula* roots (Reynoso et al., 2013), we tested whether the association of *MtNNC1* with actively translating ribosomes was affected during the root nodule symbiosis using TRAP (Translating Ribosomes Affinity Purification) in *M. truncatula* roots. TRAP is based on the expression of a FLAG-tagged version of the Ribosomal Protein of the Large Subunit 18 (FLAG-RPL18) and the affinity purification of mRNAs bound to one or more ribosomes containing FLAG-RPL18 using anti-FLAG antibodies coupled to magnetic beads followed by the isolation of the ribosome-bound RNA referred to as TRAP RNA (Zanetti et al., 2005). As a control, we also determined the association of *MtSKI3* mRNA to the ribosomes at different stages of the root nodule symbiosis. Using RT-qPCR on TRAP RNA samples, we found that the association of *MtSKI3* transcripts was upregulated not only at 2 dpi with *S. meliloti*, but also at later stages of the symbiosis, i.e., 6 and 10 dpi (Figure 4). By contrast, the association of *MtNNC1* mRNA with translating ribosomes was unaffected at 2 dpi with rhizobia but reduced at later stages, i.e. roots at 6 and 10 dpi (Figure 4), indicating that in addition to its regulation at the level of miRNA endonucleolytic cleavage, *MtNNC1* may be subjected to translational regulation, which could be mediated by miR172 associated with translating ribosomes.

**Figure 4.**
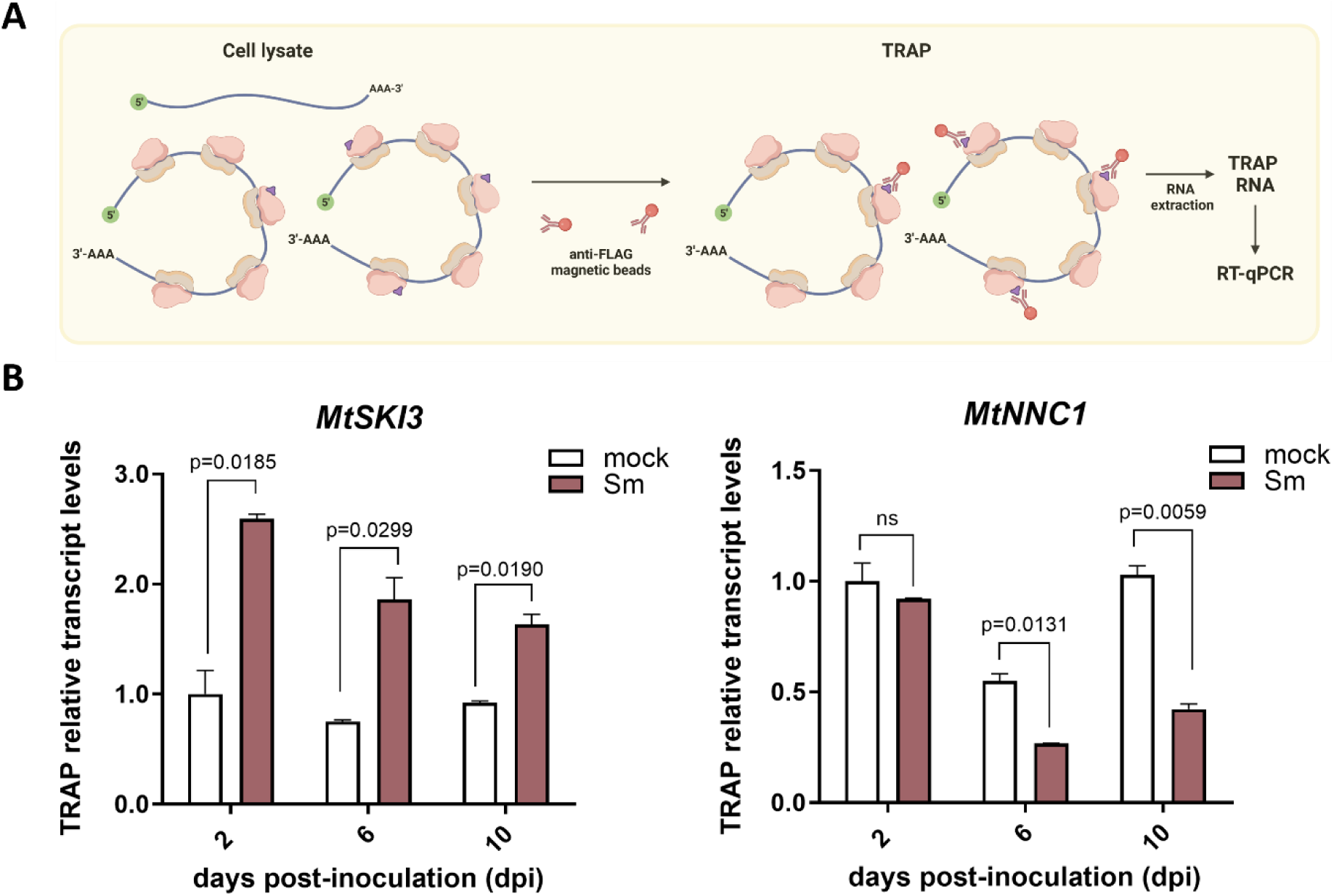
Association of *MtSKI3* and *MtNNC1* with translating ribosomes during symbiosis. **(A)** Schematic representation of the TRAP methodology. Cell lysates were prepared from root tissue expressing the FLAG-tagged MtRPL18 protein (FLAG-RPL18) under the control of the CaMV 35S promoter. mRNA bound to one or multiple ribosomes containing FLAG-tagged MtRPL18 were immunopurified using anti-FLAG coupled to magnetic beads to obtain the TRAP samples. RNA was extracted from the TRAP samples and subjected to RT-qPCR. Created with Biorender.com. **(B)** Relative transcript levels of Mt*SKI3* or *MtNNC1* in TRAP samples from roots inoculated with *S. meliloti* (Sm) or water (mock) at the indicated time points. Expression values were determined by RT-qPCR, normalized to *MtHIS3L*, and plotted relative to the mock sample at 2 dpi. Each bar represents the mean ± SEM of three biological replicates. *p* values in an unpaired two-tailed Student’s t-test are indicated for each comparison. ns: no significant differences.

### *MtSKI3* knockdown has no impact on miR172 and miR393 levels but is required for endonucleolytic cleavage

To evaluate whether the silencing of *MtSKI3* in *M. trunctula* roots affects small RNA levels and distribution, notably the miR172 and miR393 that emerged from the degradome analysis, we performed small RNA sequencing (sRNA-seq) on *GUS* RNAi and *SKI3* RNAi roots under mock conditions. No differences in the accumulation levels of miR172a, miR172b/c and miR393a/b are evident in *SKI3* RNAi roots as compared with *GUS* RNAi roots in mock conditions (Figure 5A, Supplemental Dataset 4). Stem-loop RT-qPCR confirmed sRNA-seq data, indicating that silencing of *MtSKI3* does not modify the abundance of mature miR172 or miR393 levels in *M. truncatula* roots, and further revealed that both miR172 and miR393 levels increase upon inoculation with *S. meliloti* in both *GUS* RNAi and *SKI3* RNAi roots (Figure 5B and Supplementary Figure 3). Silencing of *MtSKI3* does not lead to significantly enhanced production of 21 and 22 nt secondary siRNA derived from the *MtNNC1* or *MtAFB3* transcripts (Supplemental Figure 4) but enhances the production of 21 and 22 nt siRNAs from other loci, mainly repetitive sequences, and retrotransposon (Supplemental Figure 5, Supplemental Dataset 5). To evaluate whether there is a differential stabilization of specific miR172-directed 5′ and 3′cleavage fragments derived from the *MtNNC1* mRNA due to the silencing of *MtSKI3*, we conducted RT using random primers followed by qPCR with primer pairs designed to target the 5’ or the 3’ region of *MtNNC1* mRNA, which will detect both the full-length transcript as well the 5′or the 3′cleavage fragments, respectively. In addition, we used primers that anneal at regions flanking the miR172 cleavage site to detect exclusively the full-length *MtNNC1* mRNA (Figure 5C). This analysis revealed that the abundance of the full-length *MtNNC1* and/or the cleavage fragments do not show significant differences between SKI3 RNAi and *GUS* RNAi under mock conditions. In addition, the abundance of the full-length *MtNNC1* and/or the cleavage fragments decreases in *GUS* RNAi roots upon inoculation with *S. meliloti,* but not in *SKI3* RNAi roots (Figure 5C). These findings reveal that miR172-directed cleavage of *MtNNC1* in response to rhizobia is lost in *SKI3* RNAi roots. This effect might not be attributable to the differential stabilization of specific regions within the *MtNNC1* transcript but to other mechanisms such as alterations in RISC activity and/or mRNA degradation kinetics that underlie the observed accumulation of siRNAs derived from *MtNNC1* in *MtSKI3*-silenced roots.

**Figure 5.**
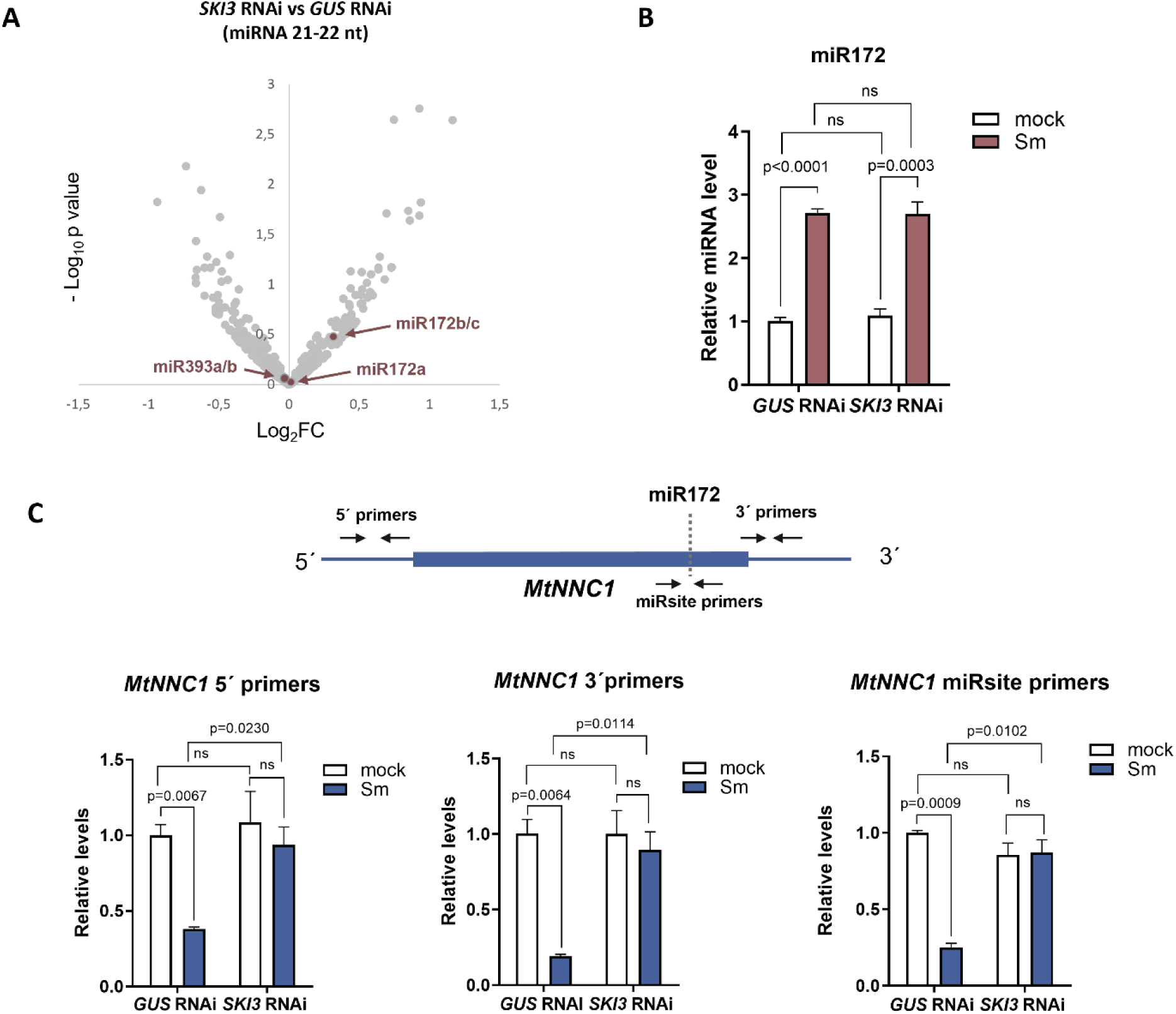
Analysis of miR172 levels and *MtNNC1* in *SKI3* RNAi and *GUS* RNAi roots. (**A**) Volcano plots showing the Log_2_FC as the mean expression level for each small RNA between *SKI3* RNAi mock and *GUS* RNAi mock samples. Each dot represents one small RNA. miR172a, miR172b/c and miR393a/b are colored black. (**B**). Stem-loop RT-qPCR of miR172 levels in *GUS* RNAi and *SKI3* RNAi roots at 2 dpi with *S. meliloti* (Sm) or water (mock). Levels were normalized to the level of miR162 and plotted relative to the mock *GUS* RNAi sample. (**C**). Relative transcript levels of *MtNNC1* in *GUS* RNAi and *SKI3* RNAi roots at 2 dpi with *S. meliloti* (Sm) or water (mock) using the primers indicated in the top panel: 5′primers, 3′primers, and miRsite primers. Expression values were determined by RT-qPCR, normalized to *MtHIS3L*, and plotted relative to the *GUS* RNAi. In (**B**) and (**C**) each bar represents the mean ± SEM of three biological replicates. *p* values of an unpaired two-tailed Student’s t-test are indicated. ns: not significant differences.

### *MtNNC1* negatively regulates nodule number and infection by *S. meliloti*

*NNC1* function has been described in determinate nodule-forming legumes, i.e. soybean and common bean (Wang et al, 2014, Nova-Franco et al, 2015). To gain insight into the function of *MtNNC1* during the formation and infection of indeterminate nodules, we used RNAi to knock down *MtNNC1* transcripts in *M. truncatula* hairy roots. This strategy decreases levels of *MtNNC1* by more than 75%b, but not the levels of its close homolog *MtERF101* (Figure 6A). The knockdown of *MtNNC1* results in a significant increase in the number of nodules formed by *S. meliloti* at all time points analyzed (7, 10, 14, and 21 dpi) compared with *GUS* RNAi roots (Figure 6B). The density of infection events also increases at 7 dpi by silencing *MtNNC1* (Figure 6C), but not their progression since the density is significantly higher only for those infection events at the microcolony stage or for infection threads (ITs) that elongate within the root hair, but not for ITs that reach the epidermal cells or the root cortex (Figure 6D). The nodulation and infection phenotypes were confirmed by a second independent RNAi construct (*MtNNC1* RNAi2), which produced similar results (Supplemental Figure 6). To better understand the molecular events affected by the knockdown of *MtNNC1* and considering that in our previous study, we observed that the induction of *MtENOD40* in response to rhizobia was impaired in *MtSKI3* silenced roots compared to control roots, we tested whether the expression of *MtENOD40* was altered by silencing of *MtNNC1*. RT-qPCR experiments revealed that *MtENOD40* transcripts accumulate to significantly higher levels when *MtNNC1* was knocked down under both mock and *S. meliloti* inoculated conditions (Figure 6E), suggesting that *MtNNC1* might act as a repressor of *MtENOD40*.

**Figure 6.**
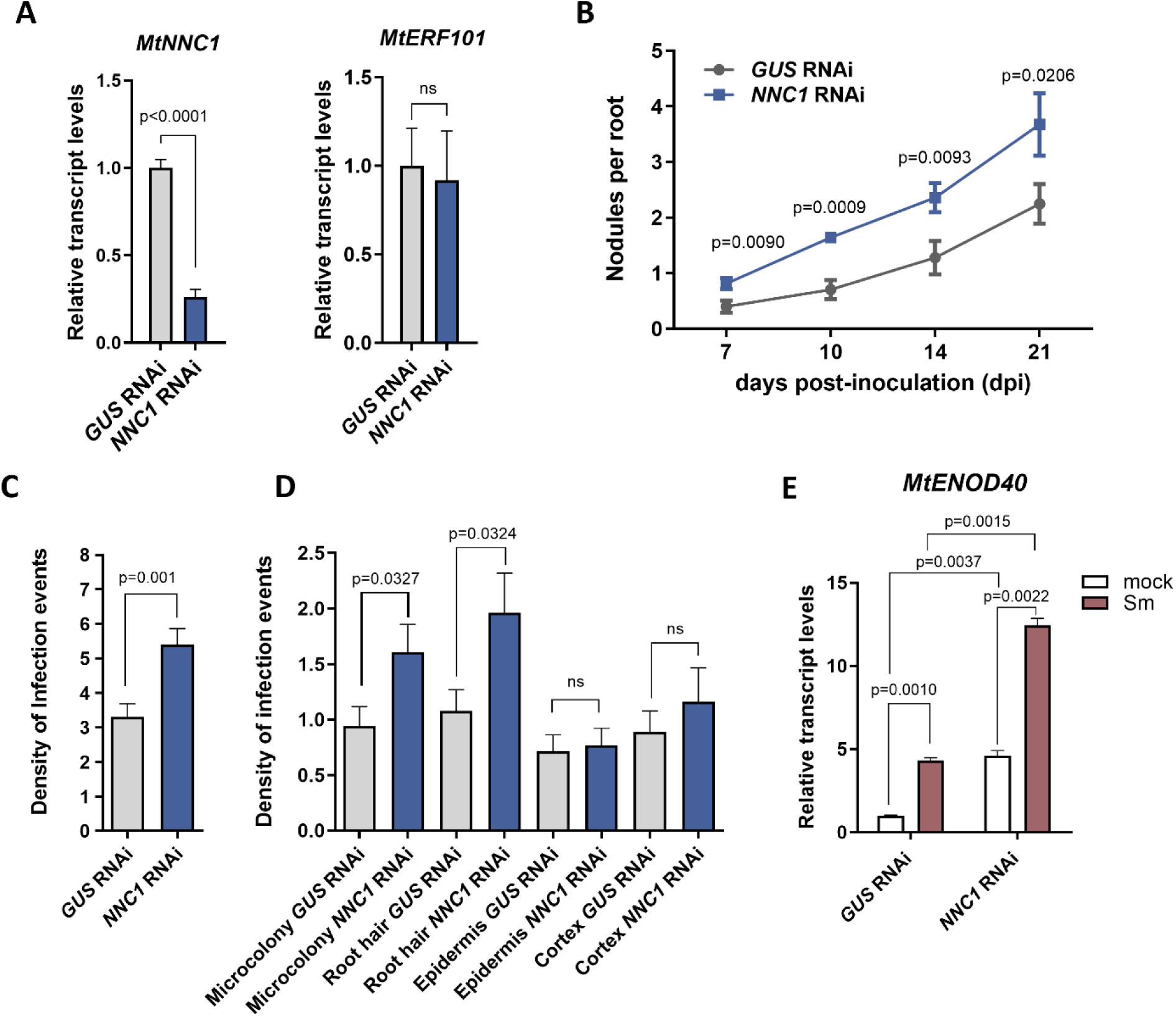
Silencing of *MtNNC1* enhanced nodulation, bacterial infection, and expression of *MtENOD40*. (**A**). Relative transcript levels of *MtNNC1* and its close homolog *MtERF101* in *GUS* RNAi and *NNC1* RNAi roots. Expression values were determined by RT-qPCR, normalized to *MtHIS3L*, and plotted relative to the level in *GUS* RNAi roots. Each bar represents the mean ± SEM of three biological replicates. *p* values in an unpaired two-tailed Student’s t-test are indicated. (**B**). Nodules per root formed in *GUS* RNAi and *NNC1* RNAi roots at 7, 10, 15, and 21 dpi with *S. meliloti*. Error bars represent the mean ± SEM of three independent biological replicates with at least 50 roots. *p* values in an unpaired two-tailed Student’s t-test between *GUS* RNAi and *MtNNC1* RNAi roots at each time point are indicated. (**C)** Infection threads (ITs) per centimeter of root developed at 7 dpi in *GUS* RNAi and *NNC1* RNAi roots. (**D**). Progression of infection events in *GUS* RNAi and *NNC1* RNAi roots. Infection events were classified as ITs that end in microcolony, in the root hair, in the epidermal cell layer, or reach the cortex at 7 dpi. In (**C**) and (**D**) each bar represents the mean ± SEM of three independent biological replicates. *p* values in an unpaired two-tailed Student’s *t*-test are indicated. (**E**). Relative transcript levels of *MtENOD40* in *GUS* RNAi and *NNC1* RNAi roots. Expression values were determined by RT-qPCR, normalized to *MtHIS3L*, and plotted relative to the *GUS* RNAi. Each bar represents the mean ± SEM of three biological replicates. *p* values in an unpaired two-tailed Student’s *t-*test in each comparison are indicated.

To investigate the functional impact of miR172 regulation on *MtNNC*1, plants ectopically overexpressing a miR172-resistant version of *MtNNC1* (r*NNC1)* were generated. RT-qPCR experiments verified that r*NNC1* expressing roots accumulated higher levels of *MtNNC1* as compared with roots transformed with the empty vector (EV) (Figure 7A). r*NNC1* roots exhibited a considerable reduction in the number of nodules formed by *S. meliloti* at all time points (7, 10, 14, and 21 dpi) analyzed as compared to the EV-expressing roots (Figure 7B). These results indicate that the overexpression of a miR172-resistant variant of *MtNNC1* negatively impacts nodule formation, suggesting that tight regulation of *MtNNC1* is crucial for the nodulation process. Indeed, *MtNNC1* knockdown shows an opposite phenotype to *SKI3* RNAi. To further dissect the regulatory dynamics between miR172 and *MtNNC1*, we analyzed the levels of miR172 transcripts in roots overexpressing *rNNC1* and EV roots at 2 dpi with *S. meliloti*. The results revealed a significant increase in miR172 accumulation in *rNNC1* roots as compared to EV roots (Figure 7C). This higher accumulation of miR172 in *rNNC1* roots exposes a regulatory feedback loop between miR172 and its target *MtNNC1*.

**Figure 7.**
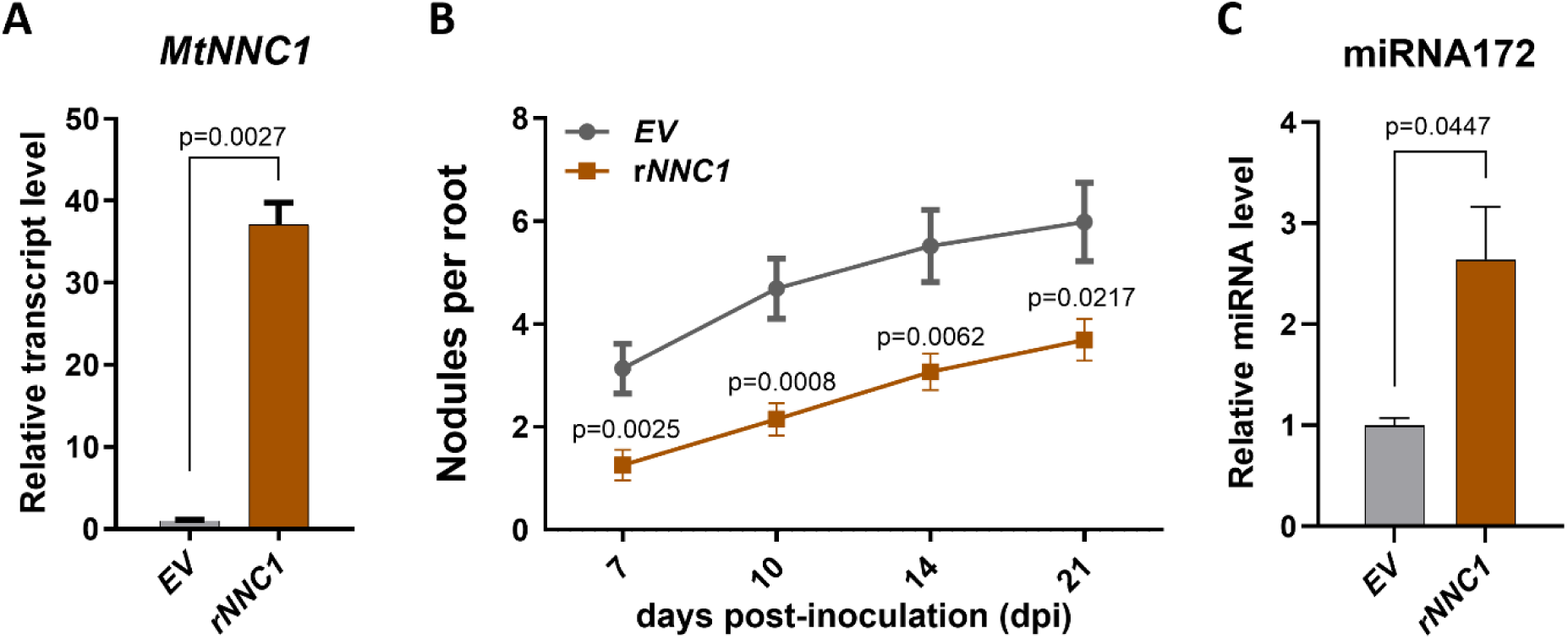
Expression of a miR172-resistant form of *MtNNC1* (*rNNC1*) reduced nodulation and increased miR172 levels. (**A**) Relative transcript levels of *MtNNC1* in EV and *rNNC1* roots. Expression values were determined by RT-qPCR, normalized to *MtHIS3L,* and plotted relative to the EV. (**B**) Nodules per root formed in EV, and *rNNC1* roots at 7, 10, 15, and 21 dpi with *S. meliloti*. Data are representative of three independent biological replicates, each with at least 60 roots. Error bars represent the mean ± SEM of three independent biological replicates. p values in an unpaired two-tailed Student’s t-test at each time point are indicated. (**C)**. Relative transcript levels of miR172 in EV, and *rNNC1* roots. Expression values were determined by stem-loop RT-qPCR, normalized to miR162a, and plotted relative to the EV. In (A) and (C) Each bar represents the mean ± SEM of three biological replicates. *p* values in an unpaired two-tailed Student’s *t*-test are indicated.

## Discussion

mRNA degradation is executed by multiple players, including 5′-to-3′exonucleases, the SKI/exosome complex that degrades deadenylated mRNAs in the 3’-to-5’ direction, and miRNAs that guide AGO proteins by base pair complementary to it targets for endonucleolytic cleavage. Here we uncover an interconnection between the SKI/exosome degradation pathway and the endonucleolytic cleavage mediated by miRNAs. Using both 5’P transcript (degradome) and sRNA-seq approaches we found that inactivation of *MtSKI3* function impairs miR172-mediated cleavage of the *MtNNC1* transcript with no alteration of miR172 levels. This suggests that the action rather than the biogenesis of miR172 is altered by the knockdown of *MtSKI3*. Upon rhizobia inoculation miR172 levels increase in both *GUS* RNAi and *SKI3* RNAi roots, in agreement with previous reports in other leguminous plants (Wang et al., 2014; Nova-Franco et al., 2015). However, whereas *MtNNC1* levels decrease upon inoculation with rhizobia in *GUS* RNAi control roots, they remain high in *SKI3* RNAi roots. This suggests that the repression of *MtNNC1* by miR172-mediated cleavage in response to rhizobia may involve *MtSKI3*. Reduced endonucleolytic cleavage was also observed for the miR393 target transcript *MtAFB3* in *SKI3* RNAi roots in mock and rhizobia inoculated conditions, despite an increase in miR393 levels upon rhizobia inoculation in both *GUS* RNAi and *SKI3* RNAi roots. Soybean TIR/AFB homologs have been shown to mediate auxin-signaling to modulate nodule number and the formation of infection foci (Cai et al., 2019). Thus, our results suggest that a functional SKI complex might be a requirement for cleavage of specific miRNA target transcripts during the root nodule symbiosis. Consistent with these results, it has been described that simultaneous mutation in genes encoding the exoribonuclease XRN4/EIN5 and SKI2 in Arabidopsis leads to overaccumulation of miRNA target transcripts, many of which encode transcription factors of the ARF, HD-ZipIII, and LBD families (Zhang et al., 2015). Besides the reduced miRNA-guided cleavage of *MtNNC1* and *MtAFB3*, we observed that neither the miR172 nor the miR393 levels changed in *MtSKI3* silenced roots as compared with control roots under mock inoculated conditions, reinforcing the idea that defects in RISC cleavage activity or the kinetic of cleavage rather than changes in miRNA levels are affected in *MtSKI3* silenced roots.

In soybean, GmNNC1 was described as a negative regulator of nodule formation and transcriptional repressor that directly binds to the *GmENOD40* promoter (Wang et al., 2014). Here, our phenotypic characterization has shown that knockdown of *MtNNC1* results in the formation of numerous nodules in *M. truncatula* roots, which agrees with that previously observed in soybean and common bean (Wang et al., 2014; Nova-Franco et al., 2015). Thus, NNC1 seems to be a negative regulator of nodulation in both determinate and indeterminate nodule-forming legumes. Moreover, levels of *MtENOD40*, which is required for nodule formation, were higher in *MtNNC1* RNAi roots under both symbiotic and non-symbiotic conditions. This is also consistent with the fact that GmNNC1 functions as a transcriptional repressor of *GmENODs* in soybean (Wang et al., 2014) as well as with increased and reduced levels of *PvENOD40* in nodules overexpressing miR172 or reduced levels in root a miR172-resistant form of the *NNC1* homolog in *P. vulgaris,* respectively (Nova-Franco et al., 2015). Remarkably, *MtNNC1* also participates in the control of rhizobia infection since silencing of *MtNNC1* results in a higher frequency of infection events, a feature that was not previously investigated in other legumes, although these infection events do not seem to be persistent since they do not progress to the root cortex. On the other hand, overexpression of a miR172-resistant variant (*rNNC1*) results in a severe reduction in nodule number in *M. truncatula*. This agrees with that previously observed in soybean and common bean (Wang et al., 2014; Nova-Franco et al., 2015), supporting the negative role of NNC1 in nodulation as a conserved feature in determinate and indeterminate nodule-forming legumes. In addition, overexpression of rNNC1 recapitulates the phenotype observed in *MtSKI3* silenced roots (Traubenik et al., 2020), reinforcing the idea that *MtSKI3* is required for proper downregulation of *MtNNC1* during symbiosis. Interestingly, the expression of a miR172-resistant version of *MtNNC1* (*rNNC1*) leads to elevated miR172 levels, suggesting a potential compensatory mechanism that counteracts the overexpression of *rNNC1* to restore the balance in the miR172-*MtNNC1* regulatory axis. This differs from that previously described in soybean, where miR172 levels were reduced in roots overexpressing a miR172-resistant variant of *GmNNC1*. Moreover, in soybean, miR172 is transcriptionally repressed by *GmNNC1* (Wang et al., 2019). The opposite behavior of *M. truncatula* and soybean might reflect differences in the regulatory mechanisms for miR172 transcriptional activation between the two legume species, i.e. *M. truncatula* and soybean, but also differences in the time points after rhizobia inoculation used to evaluate miR172 levels.

Endonucleolytic cleavage of mRNAs mediated by miRNAs and/or siRNAs, as well as the production of siRNAs, was proposed to occur in polysomes (Brodersen et al., 2008; Liu et al., 2012; Li et al., 2016; Traubenik et al., 2020). Moreover, miRNA-mediated translational inhibition of target mRNAs seems to be a widespread mechanism present in Arabidopsis and maize plants (Brodersen et al., 2008; Yang and Thompson, 2024). The miR172 is one of the miRNAs that, in addition to acting at the endonucleolytic cleavage level (Schwab et al., 2005), has been shown to repress its target, the homeotic gene AP2, through translation inhibition in Arabidopsis (Aukerman and Sakai, 2003; Chen, 2004). We previously demonstrated that miR172 is highly associated with polysomes in *M. truncatula* roots (Reynoso et al., 2013). Here, we found that *MtNNC1* was associated with polysomes and that its association with polysomes decreased in roots in response to inoculation with *S. meliloti* at 6 and 10 dpi, while *MtSKI3* increased its association with polysomes as early as 2 dpi and remained high up to 10 dpi. This indicates that regulation of *MtSKI3* at the translational level could impact the miR172-mediated endonucleolytic cleavage, which most likely occurs while *MtNNC1* is bound to the polysomes, although translational inhibition and stalling of *MtNNC1* during symbiosis cannot be excluded. The SKI complex has been found bound to ribosomes in mammals and yeast, likely facilitating the delivery of 80S ribosome-bound translationally stalled mRNAs to the exosome for mRNA decay (Kögel et al., 2022). The requirement of *MtSKI3* observed here for miR172-mediated repression of *MtNNC1,* as well as for other miRNA targets (e.g. *AFB3*), might reflect a function of the plant SKI complex to extract translational repressed mRNAs and/or the 5′products of miRNA cleavage and their delivery to the exosome for 3′to 5′degradation. This study highlights the fine-tuning of gene regulation involving miRNAs, translation, and mRNA decay processes required for the establishment of a successful symbiosis between legume plants and nitrogen-fixing bacteria.

## Material and Methods

### Biological material and vector construction

*Medicago truncatula* Jemalong A17 seeds were obtained from INRA Montpellier, France (http://www.montpellier.inra.fr). *SKI3* RNAi and GUS RNAi constructs were previously generated (Traubenik et al, 2020). The *MtNNC1* RNAi 1 or *MtNNC1* RNAi 2 constructs were generated by amplification of *MtNNC1* fragment using *M. truncatula* cDNA as a template and the MtNNC1RNAi 1 F and MtNNC1RNAi 1R or MtNNC1RNAi 2 F and MtNNC1RNAi 2 R primers, respectively, wich are listed in Supplemental Dataset 6. The *MtNNC1* amplified fragments were cloned into the pTOPO/ENTR vector (Thermo Fisher Scientific) and recombined into the destination vector pK7GWIWG2DII (Karimi et al., 2007). The miR172-resistant variant of *MtNNC1* (mirResMtNNC1) was generated by PCR-directed mutagenesis, introducing specific nucleotide changes (CAT to GCA) at the 9^th^, 10^th^, and 11^th^ positions of the miR172 binding site while preserving the encoded serine residues. These modifications were made using the mirResMtNNC1 F and mirResMtNNC1 R primers listed in Supplemental Dataset 6. The amplified fragment was cloned into the pENTR/D-TOPO vector and then recombined into the Gateway-compatible binary vector pK7WG2D,1 (Karimi et al., 2002), in which the expression of *MtNNC1* is driven by the cauliflower mosaic virus *35S* promoter. All constructs were verified by sequencing. Binary vectors were introduced into *A. rhizogenes* Arqua1 (Quandt et al., 1993) by electroporation. *Sinorhizobium meliloti* strain 1021 (Meade and Signer, 1977) or the same strain expressing RFP (Tian et al., 2012) were used for root inoculation as previously described (Hobecker et al., 2017).

### Plant growth conditions, hairy root transformation, and rhizobia inoculation

Seeds were sterilized and germinated as previously described (Traubenik et al, 2020). Germinated seedlings were transferred to Petri dishes containing agar Fahraeus media (Fahraeus, 1957) covered with sterile filter paper. Transgenic roots were generated by *Agrobacterium rhizogenes*-mediated transformation as previously described (Boisson-Dernier et al., 2001). Plants that developed hairy roots were transferred to slanted boxes containing Fahraeus media free of nitrogen. Seedlings were grown at 25°C with a long day period (16-h day/8-h night cycle) and 50% humidity. Roots were inoculated with 10 ml of a 1:1000 dilution of *S. meliloti* 1021 (Meade and Signer, 1977) culture grown in liquid TY media until OD_600_ reached 0.8 or with 10 ml of water as a control (mock treatment). One hour later, the excess liquid was discarded, and seedlings were incubated vertically under the temperature and light conditions for growth described above. For RNA isolation, root tissue was harvested, frozen in liquid N_2,_ and stored at -80°C.

### GMUCT 2.0 and Small RNA libraries preparation

GMUCT 2.0 and Small RNA libraries were prepared from RNA samples isolated from *GUS* RNAi or *SKI3* RNAi roots inoculated with water (mock) or *S. meliloti* for 48 hours. Whole root tissue was collected from more than 300 plants and pooled. Three biological replicates were conducted consisting of independent experiments performed on different days. GMUCT 2.0 libraries were constructed following Willmann et al., 2014 replacing kit components with commercially available supplies. Briefly, RNA was extracted using Trizol (Thermo Scientific) and 30 µg of each sample was used to obtain PolyA RNA using oligo dT conjugated to magnetic beads. The 5′adapter was ligated to the RNA using T4RNA ligase 1 and a second round of polyA selection was performed in order to purify unligated 5′adapters. The selected RNA was ligated to a 3’ adapter and after reverse transcription, the library was amplified with PCR primers from BrAD-seq (Townsley *et. al.*, 2015). KAPA HiFi DNA Polymerase (Kapa Biosystems) was used for amplification. Libraries were characterized using gel electrophoresis and the amplified libraries were isolated and sequenced using the Illumina NextSeq500 platform at the IIGB Genomics Core facility at UC Riverside (https://genomics.iigb.ucr.edu/).

Small RNA libraries were constructed using the NEBNext® Multiplex Small RNA Library Prep Set for Illumina following the manufacturer’s instructions, starting with 500 ng of Total RNA extracted with Trizol. Briefly, 3′ and 5’ adaptors were ligated to the ends of the single-stranded RNA. After ligation of the adaptors, the small RNAs were used as a template for the synthesis of one complementary strand of DNA. Reverse transcribed molecules that contain both adaptor sequences were then amplified by PCR. Libraries were characterized using gel electrophoresis and the amplified libraries were isolated. Small RNA-seq libraries were sequenced using the Illumina HiSeq 2500 RP platform at BGI Korea (https://www.bgi.com/global).

The size of DNA fragments in the GMUCT 2.0 and Small RNA libraries was verified on an Agilent 2100 Bioanalyzer using the DNA-HS kit (Agilent).

### GMUCT2-seq analysis

GMUCT2.0 reads were aligned to the *M. truncatula* A17 Mt5.0 mRNAs (Pecrix et al., 2018) in the CyVerse Discovery Environment using STAR v2.4.0.1 reporting the best alignment with ≤2 nt mismatches. Count abundances were calculated using eXpress v1.5.1, with default parameters. Peaks were detected with the “Findpeaks” function of the HOMER package (Heinz et al., 2010) with the parameters “-regionRes 1”, “-minDist 150” and “-region”. Peaks overlapping in at least two replicates were conserved for further analysis using the function “findOverlapsOfPeaks” of the ChipPeakAnno R package (Zhu et al, 2010). Reads in the peaks were quantified in each replicate using the “summarizeOverlaps” function of the GenomicRanges R package (Lawrence et). miRNAs sequences were obtained from miRbase (v22) (Kozomara and Griffiths-Jones, 2014)) and target sites were predicted using Miranda software (Enright et al.,2003) over cDNA sequences of Mt5.0 genome with parameters of gap-opening penalty set to -2 and energy threshold of -22 kcal/mol. Genomic locations predicted for 3’ fragments resulting from miRNA cut-site were deducted. Reads were quantified strictly over those genomic coordinates. The counts for peaks and cut sites were statistically evaluated using DESeq2 (Love et al., 2014). Genotype differences over features that had a log fold change value of 1 or more, or -1 or less, and a p-value < 0.05 were identified as differential GMUCT regions or cut-sites.

### Small RNA-seq analysis

Following sequencing, small RNA sequencing adapters were removed using cutadapt (v2.10). Sequences matching ribosomal or transfer RNA were filtered out using bowtie (v1.3.1). Reads of 21 and 22 nucleotides were mapped on the Mt5.0 genome (Pecrix et al., 2018) with ShortStack (v3.8.5) under strict no-mismatch conditions (--mismatches 0), keeping all primary multi-mapping (--bowtie_m all) and correcting for multi-mapped reads based on uniquely mapped reads (--mmap u). The read accumulation of 21/22nt for each gene annotation was quantified using ShortStack (Axtell, 2013). To assess miRNA abundance, the count of sequences corresponding to each mature miRNA in *Medicago* miRBase (v22) was used (Kozomara and Griffiths-Jones, 2014). Differential 21/22nt sRNA accumulation and log2 fold changes between conditions were computed using DESeq2 (v1.34.0) (Love et al., 2014). FDR correction of the p-value was used.

### Isolation of polysomes by TRAP

Isolation of polysomes by TRAP was accomplished as previously described (Traubenik et al., 2020). TRAP material was subjected to RNA extraction using Trizol following the manufacturer’s recommendations (Thermo Scientific).

### RT-qPCR

Total RNA was extracted using Trizol (Thermo Fisher Scientific), following the manufacturer’s recommendations, and digested with RNase-free DNase (Promega). Total and TRAP RNA samples were subjected to first-strand cDNA synthesis using Moloney Murine Leukemia Virus reverse transcriptase (Promega). Expression analysis by RT-qPCR was performed using the iQ SYBR Green Supermix kit (Bio-Rad) and the CFX96 qPCR system (Bio-Rad) as described previously (Blanco et al., 2009). For each pair of primers, the presence of a unique PCR product of the expected size was verified in agarose gels. The *M. truncatula HISTONE LIKE 3 (HIS3L-* MtrunA17_Chr4g0054151) was selected as a reference transcript for the normalization of RT-qPCR data based on previously reported geNORM analysis (Reynoso et al., 2013). Primers used are listed in Supplemental Dataset 6. microRNA quantification was performed by stem-loop RT-qPCR as described previously (Hobecker et al., 2017) using the primers listed in Supplemental Dataset 6. miR162 was used as a reference transcript for normalization.

### Phenotypic Analysis

Nodule number was recorded at different time points after inoculation with *S. meliloti* as described previously by Hobecker et al. (2017). Infection events were quantified at 7 dpi as described by Traubenik et al. (2020). The statistical significance of the differences for each parameter was determined by unpaired two-tailed Student’s *t-tests* for each construct.

### Accession Numbers

Raw sequence files (fastq files) supporting the conclusions of this article were deposited at Gene Expression Omnibus under series entry numbers GEO GSE135920 and GSE285134. All processed data were included in Supplemental Datasets 1-5.

Sequence data from this article can be found in MtrunA17r5.0-ANR, (https://medicago.toulouse.inra.fr/MtrunA17r5.0-ANR/) or in miRbase (v22) (http://www.mirbase.org) under the following accession numbers: *MtSKI3*<colcnt=6> (MtrunA17_Chr5g0393431), *MtNNC1* (MtrunA17_Chr4g0030191), *MtENOD40* (MtrunA17_Chr8g0368441), *MtHISL3* (MtrunA17_Chr4g0054151), *MtERF1* (MtrunA17_Chr2g0325911), miR172a (MI0005600), miR172b (MI0018368), miR393a (MI0001745), miR393b (MI0005601), miR162 (MI0001738).

## Acknowledgments

This work was supported by grants of the Agencia Nacional de Promoción de la Investigación, el Desarrollo Tecnológico y la Innovación (Agencia I+D+I) of Argentina, FONCYT (PICT2019-00554, PICT2019-0029, PICT2020-00053, and PICT-2021-I-A-00170), the Ministerio de Ciencia, Tecnología e Innovación (MINCyT) of Argentina (RIBOLEG, CONVE-2023-100766842), the Centre National de la Recherche Scientifique (CNRS) through the International Research Project LOCOSYM, and Saclay Plant Sciences-SPS (ANR-17-EUR-0007). S.T. is supported by the LUMIROOT (101110703) project funded by the Marie Skłodowska-Curie Actions and the MICROLUP project (ANR MICROLUP 19-CE13-0029-02NA) funded by the Agence Nationale de la Recherche ANR, and was funded by a CONICET fellowship and a Fulbright-Williams Foundation fellowship. M.A.R., F.A.B., and M.E.Z. are members of CONICET, Argentina. F.S-R. is funded by a Saclay Plant Sciences-SPS fellowship. M.Y. is funded by a CONICET fellowship. A.C., T.B., and M.C. are members of CNRS, France. J.B-S. is a member of NSF, USA.

## Author Contributions

S.T., M.E.Z., F.B., M.C., and J.B-S. designed the research. S.T., M.A.R., F.S-R, M.Y., M.H., A.C., and T.B. performed the research. S.T. and M.E.Z. wrote the original draft of the article. S.T., M.A.R., F.S-R., M.Y., T.B., and M.E.Z. analyzed the data. S.T., M.A.R., A.C., T.B., M.C., J.B-S., F.B., and M.E.Z. reviewed and edited the article. Funding was acquired by M.E.Z., F.B., M. C., J.B-S, and S.T. M.E.Z., F.B., M.C., T.B., and J.B-S. supervised the study.

## References

1. Allen E, Xie Z, Gustafson AM, Carrington JC (2005) microRNA-directed phasing during trans-acting siRNA biogenesis in plants. Cell 121: 207–221

2. Arribere JA, Fire AZ (2018) Nonsense mRNA suppression via nonstop decay. Elife 7

3. Aukerman MJ, Sakai H (2003) Regulation of flowering time and floral organ identity by a MicroRNA and its APETALA2-like target genes. Plant Cell 15: 2730–2741

4. Axtell MJ (2013) ShortStack: comprehensive annotation and quantification of small RNA genes. Rna 19: 740–751

5. Bartel DP (2004) MicroRNAs: genomics, biogenesis, mechanism, and function. Cell 116: 281–297

6. Betel D, Koppal A, Agius P, Sander C, Leslie C (2010) Comprehensive modeling of microRNA targets predicts functional non-conserved and non-canonical sites. Genome Biol 11: R90

7. Branscheid A, Marchais A, Schott G, Lange H, Gagliardi D, Andersen SU, Voinnet O, Brodersen P (2015) SKI2 mediates degradation of RISC 5’-cleavage fragments and prevents secondary siRNA production from miRNA targets in Arabidopsis. Nucleic Acids Res 43: 10975–10988

8. Brodersen P, Sakvarelidze-Achard L, Bruun-Rasmussen M, Dunoyer P, Yamamoto YY, Sieburth L, Voinnet O (2008) Widespread Translational Inhibition by Plant miRNAs and siRNAs. Science 320: 1185–1190

9. Brodersen P, Sakvarelidze-Achard L, Bruun-Rasmussen M, Dunoyer P, Yamamoto YY, Sieburth L, Voinnet O (2008) Widespread translational inhibition by plant miRNAs and siRNAs. Science 320: 1185–1190

10. Cai Z, Wang Y, Zhu L, Tian Y, Chen L, Sun Z, Ullah I, Li X (2017) GmTIR1/GmAFB3-based auxin perception regulated by miR393 modulates soybean nodulation. New Phytol 215: 672–686

11. Cai Z, Zeng DE, Liao J, Cheng C, Sahito ZA, Xiang M, Fu M, Chen Y, Wang D (2019) Genome-Wide Analysis of Auxin Receptor Family Genes in Brassica juncea var. tumida. Genes (Basel) 10

12. Carpentier M-C, Deragon J-M, Jean V, Be SHV, Bousquet-Antonelli C, Merret R (2020) Monitoring of XRN4 Targets Reveals the Importance of Cotranslational Decay during Arabidopsis Development Plant Physiology 184: 1251–1262

13. Chantarachot T, Bailey-Serres J (2018) Polysomes, Stress Granules, and Processing Bodies: A Dynamic Triumvirate Controlling Cytoplasmic mRNA Fate and Function. Plant Physiol 176: 254–269

14. Chen X (2004) A microRNA as a translational repressor of APETALA2 in Arabidopsis flower development. Science 303: 2022–2025

15. Halbach F, Reichelt P, Rode M, Conti E (2013) The yeast ski complex: crystal structure and RNA channeling to the exosome complex. Cell 154: 814–826

16. Hou CY, Lee WC, Chou HC, Chen AP, Chou SJ, Chen HM (2016) Global Analysis of Truncated RNA Ends Reveals New Insights into Ribosome Stalling in Plants. Plant Cell 28: 2398–2416

17. Kögel A, Keidel A, Bonneau F, Schäfer IB, Conti E (2022) The human SKI complex regulates channeling of ribosome-bound RNA to the exosome via an intrinsic gatekeeping mechanism. Mol Cell 82: 756–769.e758

18. Kozomara A, Griffiths-Jones S (2014) miRBase: annotating high confidence microRNAs using deep sequencing data. Nucleic Acids Res 42: D68–73

19. Lange H, Gagliardi D (2021) Catalytic activities, molecular connections, and biological functions of plant RNA exosome complexes. The Plant Cell 34: 967–988

20. Li S, Le B, Ma X, You C, Yu Y, Zhang B, Liu L, Gao L, Shi T, Zhao Y, Mo B, Cao X, Chen X (2016) Biogenesis of phased siRNAs on membrane-bound polysomes in Arabidopsis. Elife 5

21. Liu L, Chen X (2016) RNA Quality Control as a Key to Suppressing RNA Silencing of Endogenous Genes in Plants. Mol Plant 9: 826–836

22. Liu MJ, Wu SH, Chen HM (2012) Widespread translational control contributes to the regulation of Arabidopsis photomorphogenesis. Mol Syst Biol 8: 566

23. Love MI, Huber W, Anders S (2014) Moderated estimation of fold change and dispersion for RNA-seq data with DESeq2. Genome Biology 15: 550

24. Manavella PA, Hagmann J, Ott F, Laubinger S, Franz M, Macek B, Weigel D (2012) Fast-forward genetics identifies plant CPL phosphatases as regulators of miRNA processing factor HYL1. Cell 151: 859–870

25. Nova-Franco B, Iniguez LP, Valdes-Lopez O, Alvarado-Affantranger X, Leija A, Fuentes SI, Ramirez M, Paul S, Reyes JL, Girard L, Hernandez G (2015) The micro-RNA72c-APETALA2-1 node as a key regulator of the common bean-Rhizobium etli nitrogen fixation symbiosis. Plant Physiol 168: 273–291

26. Orban TI, Izaurralde E (2005) Decay of mRNAs targeted by RISC requires XRN1, the Ski complex, and the exosome. Rna 11: 459–469

27. Pelechano V, Wei W, Steinmetz LM (2015) Widespread Co-translational RNA Decay Reveals Ribosome Dynamics. Cell 161: 1400–1412

28. Potuschak T, Vansiri A, Binder BM, Lechner E, Vierstra RD, Genschik P (2006) The exoribonuclease XRN4 is a component of the ethylene response pathway in Arabidopsis. Plant Cell 18: 3047–3057

29. Reynoso MA, Blanco FA, Bailey-Serres J, Crespi M, Zanetti ME (2013) Selective recruitment of mRNAs and miRNAs to polyribosomes in response to rhizobia infection in Medicago truncatula. Plant J 73 289–301

30. Rymarquis LA, Souret FF, Green PJ (2011) Evidence that XRN4, an Arabidopsis homolog of exoribonuclease XRN1, preferentially impacts transcripts with certain sequences or in particular functional categories. RNA 17: 501–511

31. Schmidt C, Kowalinski E, Shanmuganathan V, Defenouillère Q, Braunger K, Heuer A, Pech M, Namane A, Berninghausen O, Fromont-Racine M, Jacquier A, Conti E, Becker T, Beckmann R (2016) The cryo-EM structure of a ribosome-Ski2-Ski3-Ski8 helicase complex. Science 354: 1431–1433

32. Schwab R, Palatnik JF, Riester M, Schommer C, Schmid M, Weigel D (2005) Specific effects of microRNAs on the plant transcriptome. Dev Cell 8: 517–527

33. Szádeczky-Kardoss I, Csorba T, Auber A, Schamberger A, Nyikó T, Taller J, Orbán TI, Burgyán J, Silhavy D (2018) The nonstop decay and the RNA silencing systems operate cooperatively in plants. Nucleic Acids Res 46: 4632–4648

34. Traubenik S, Reynoso MA, Hobecker K, Lancia M, Hummel M, Rosen B, Town C, Bailey-Serres J, Blanco F, Zanetti ME (2020) Reprogramming of Root Cells during Nitrogen-Fixing Symbiosis Involves Dynamic Polysome Association of Coding and Noncoding RNAs. The Plant Cell 32: 352–373

35. Vaucheret H (2006) Post-transcriptional small RNA pathways in plants: mechanisms and regulations. Genes Dev 20: 759–771

36. Vigh ML, Bressendorff S, Thieffry A, Arribas-Hernández L, Brodersen P (2022) Nuclear and cytoplasmic RNA exosomes and PELOTA1 prevent miRNA-induced secondary siRNA production in Arabidopsis. Nucleic Acids Res 50: 1396–1415

37. Wang L, Sun Z, Su C, Wang Y, Yan Q, Chen J, Ott T, Li X (2019) A GmNINa-miR172c-NNC1 Regulatory Network Coordinates the Nodulation and Autoregulation of Nodulation Pathways in Soybean. Mol Plant 12: 1211–1226

38. Wang Y, Wang L, Zou Y, Chen L, Cai Z, Zhang S, Zhao F, Tian Y, Jiang Q, Ferguson BJ, Gresshoff PM, Li X (2014) Soybean miR172c targets the repressive AP2 transcription factor NNC1 to activate ENOD40 expression and regulate nodule initiation. Plant Cell 26: 4782–4801

39. Willmann MR, Berkowitz ND, Gregory BD (2014) Improved genome-wide mapping of uncapped and cleaved transcripts in eukaryotes--GMUCT 2.0. Methods 67: 64–73

40. Yang H, Thompson B (2024) Widespread changes to the translational landscape in a maize microRNA biogenesis mutant. The Plant Journal 119: 1986–2000

41. Yoshikawa M, Peragine A, Park MY, Poethig RS (2005) A pathway for the biogenesis of trans-acting siRNAs in Arabidopsis. Genes Dev 19: 2164–2175

42. Yu A, Saudemont B, Bouteiller N, Elvira-Matelot E, Lepere G, Parent JS, Morel JB, Cao J, Elmayan T, Vaucheret H (2015) Second-Site Mutagenesis of a Hypomorphic argonaute1 Allele Identifies SUPERKILLER3 as an Endogenous Suppressor of Transgene Posttranscriptional Gene Silencing. Plant Physiol 169: 1266–1274

43. Yu X, Willmann MR, Anderson SJ, Gregory BD (2016) Genome-Wide Mapping of Uncapped and Cleaved Transcripts Reveals a Role for the Nuclear mRNA Cap-Binding Complex in Cotranslational RNA Decay in Arabidopsis. Plant Cell 28: 2385–2397

44. Zanetti ME, Chang IF, Gong F, Galbraith DW, Bailey-Serres J (2005) Immunopurification of polyribosomal complexes of Arabidopsis for global analysis of gene expression. Plant Physiol 138: 624–635

45. Zhang X, Zhu Y, Liu X, Hong X, Xu Y, Zhu P, Shen Y, Wu H, Ji Y, Wen X, Zhang C, Zhao Q, Wang Y, Lu J, Guo H (2015) Plant biology. Suppression of endogenous gene silencing by bidirectional cytoplasmic RNA decay in Arabidopsis. Science 348: 120–123

46. Zhao L, Kunst L (2016) SUPERKILLER Complex Components Are Required for the RNA Exosome-Mediated Control of Cuticular Wax Biosynthesis in Arabidopsis Inflorescence Stems. Plant Physiol 171: 960–973

47. Zinoviev A, Ayupov RK, Abaeva IS, Hellen CUT, Pestova TV (2020) Extraction of mRNA from Stalled Ribosomes by the Ski Complex. Mol Cell 77: 1340–1349.e1346

